# TNF-α differentially regulates cell cycle genes in promyelocytic and granulocytic HL-60/S4 cells

**DOI:** 10.1101/471789

**Authors:** Elsie C. Jacobson, Lekha Jain, Jo K. Perry, Mark H. Vickers, Ada L. Olins, Donald E. Olins, Justin M. O’Sullivan

**Author notes:** Author email addresses: ECJ < >, LJ, JKP < >, ALO < >, DEO < >, MHV < >, JMO < >.

## Abstract

Tumor necrosis factor alpha (TNF-α) is a potent cytokine involved in systemic inflammation and immune modulation. Signaling responses that involve TNF-α are context dependent and capable of stimulating pathways promoting both cell death and survival. TNF-α treatment has been investigated as part of a combined therapy for acute myeloid leukemia due to its modifying effects on all-trans retinoic acid (ATRA) mediated differentiation into granulocytes.

To investigate the interaction between cellular differentiation and TNF-α, we performed RNA-sequencing on two forms of the human HL-60/S4 promyelocytic leukemia cell line treated with TNF-α. The ATRA-differentiated granulocytic form of HL-60/S4 cells had an enhanced transcriptional response to TNF-α treatment compared to the undifferentiated promyelocytes. The observed TNF-α responses included differential expression of cell cycle gene sets, which were generally upregulated in TNF-α treated promyelocytes, and downregulated in TNF-α treated granulocytes. This is consistent with TNF-α induced cell cycle repression in granulocytes and cell cycle progression in promyelocytes. Moreover, comparisons with gene expression changes associated with differentiation indicated that TNF-α treatment of granulocytes shifts the transcriptome towards that of a macrophage.

We conclude that TNF-α treatment promotes a divergent transcriptional program in promyelocytes and granulocytes. TNF-α promotes cell cycle associated gene expression in promyelocytes. In contrast, TNF-α stimulated granulocytes have reduced cell cycle gene expression, and a macrophage-like transcriptional program.

## 1. Background

Tumor necrosis factor alpha (TNF-α) is a pro-inflammatory cytotoxic cytokine [1] which activates the innate immune response [2–4], and induces migration [5,6] and production of pro-inflammatory cytokines [7,8]. Dysregulation of TNF-α can be a factor in autoimmune disease [9,10] and anti-TNF antibodies are used to treat a range of inflammatory disorders [11–13]. Initially investigated as a cancer therapeutic due to its ability to promote apoptotic cell death specifically of tumor cells [14], systemic TNF-α treatment has failed clinical trials as a solo cancer therapeutic as it induces unacceptable levels of toxicity [15].

TNF-α signaling is complex with numerous and sometimes conflicting responses being modulated by interaction with two cell surface TNF-α receptors, TNFR1 and TNFR2 [1]. TNF-α binding can have a wide range of effects via activation of signal transduction pathways, including all three groups of mitogen activated kinases (MAPK); extracellular-signal-regulated kinases (ERKs), the cJun NH2-terminal kinases (JNKs), and the p38 MAP kinases [16], which each have complex regulatory effects on the cellular phenotype [17,18]. TNF-α signaling leads to transcriptional upregulation of pro-inflammatory cytokines including *IL-6* [7] and *TNF* itself [8], resulting in pro-inflammatory feedback loops [19]. Notably, TNFR1 and TNFR2 have individual and combinatorial effects on cell death and inflammation [1,20,21]. TNFR1 signaling induces pro-apoptotic pathways resulting in caspase activation, and pro-survival Nuclear Factor Kappa B (NFKB) signaling [22,23]. For example, hematopoietic cells growing in log phase rapidly undergo apoptosis in response to TNF, while quiescent cells in stationary phase re-enter the cell cycle on TNF-α stimulation [24]. These apparently conflicting TNF-α responses can be explained by temporal and developmental effects that include cell type [25], receptor expression [24], priming with cytokines or inflammatory stimuli [26,27], and cell cycle stage [28].

The HL-60/S4 cell line was derived from an acute promyelocytic leukemia patient [29]. These promyelocytic cells can be differentiated into granulocytic or macrophage forms with the addition of all-trans retinoic acid (ATRA) or 12-O-tetradecanoylphorbol-13-acetate (TPA), respectively [30]. Differentiation into the granulocytic form slows cell growth [30] and ultimately leads to cell death [31]. This discovery lead to the clinical use of ATRA as a treatment for acute promyeloid leukemia [32]. Combined treatment with ATRA and TNF-α enhances differentiation of myelogenous leukemia cells, and therefore has been investigated as a synergistic therapy [33,34]. Notably, ATRA-induced differentiation activates components of the TNF-α signaling pathway [33].

A previous study demonstrated differential effects of TNF-α treatment on candidate genes in HL-60 cells before and after ATRA treatment [35]. Here, we investigate the genome-wide transcriptional response to TNF-α treatment of the promyelocytic and granulocytic forms of HL-60/S4 cells. We identify a conserved inflammatory and apoptotic response to TNF-α treatment in both promyelocytic and granulocytic cells. We also identify opposing effects of TNF-α treatment on the expression of cell cycle genes, supporting cell cycle progression in promyelocytes and cell cycle repression in granulocytes. We propose that the different TNF-α mediated responses arise through sets of genes being responsive to different thresholds of total (endogenous and exogenous) TNF-α levels.

## 2. Results

In order to investigate transcriptional responses to TNF-α, the promyelocytic cell line HL-60/S4 was differentiated into a granulocytic form by treatment with ATRA for 96 hours, as described previously [36]. Both undifferentiated (promyelocytic) and differentiated (granulocytic) cells were treated with 16ng/mL TNF-α in calcium supplemented media for two hours. TNF-α treatment of promyelocytes and granulocytes were run as independent experiments; as a consequence, analysis of transcriptional changes following differentiation was not appropriate. However, a recently published study investigated transcriptional changes following differentiation of promyelocytic HL-60/S4 cells into granulocytes or macrophages [30]. We reanalyzed this data for use in comparisons with the TNF-α response. All genes in both sets of data that were significantly differentially expressed (False discovery rate adjusted p-value (FDR)<0.05) following differentiation or TNF-α treatment were identified (Supp table 1). In order to assess the reproducibility of our data, a subset of differentially expressed genes (*TNF, CDK1, VCAM1*) were analyzed by RT-qPCR qPCR in a set of independent samples treated as above *i.e*. ± 1 μM ATRA and ± 16ng/mL TNF-α (Supp table 2). Consistent with RNA-seq analysis, RT-qPCR demonstrated that TNF-α stimulated promyelocytes and granulocytes have increased levels of *TNF* and *VCAM1*. *CDK1* increases expression in promyelocytes and decreases expression in granulocytes following TNF-α treatment.

### 2.1 Gene expression changes after TNF-α treatment of promyelocytic and granulocytic cells

RNA-seq libraries were aligned using an ultrafast universal RNA-seq aligner (STAR [37]) and assigned to genes with featureCounts [38]. DESeq2 [39] was used to filter out lowly-expressed genes and performing differential expression analysis. 14,420 genes were analyzed for TNF-α dependent differential expression in promyelocytes. 21,305 genes were analyzed for TNF-α dependent differential expression in granulocytes. Principal component analysis of variance stabilized transcripts confirmed clustering by treatment (Supp fig 1).

The promyelocytic and granulocytic forms of HL-60/S4 cells both exhibited strong transcriptional responses to TNF-α treatment (Supp table 1). In promyelocytes, 1,312 genes were significantly increased and 980 significantly decreased (FDR adjusted p value < 0.05, Fig 1A). TNF-α treatment of granulocytes significantly increased expression of 3809 genes and decreased expression of 3,597 genes (FDR adjusted p value < 0.05, Fig 1B). Notably, the granulocytic form had more than three times the number of significantly differentially expressed genes (Fig 1C). Despite this, there was significant overlap in the TNF-α dependent transcriptional response between the granulocytic and promyelocytic cells (2.5 fold more than expected by chance, p<1×10^−5^, bootstrapping). The TNF-α treatment consistently resulted in more genes being upregulated than downregulated in both the promyelocytic and granulocytic forms of the HL-60/S4 cells (p<2×10^−16^ and p=5×10^−4^, respectively; 2-sample test for equality of proportions).

**Figure 1.**
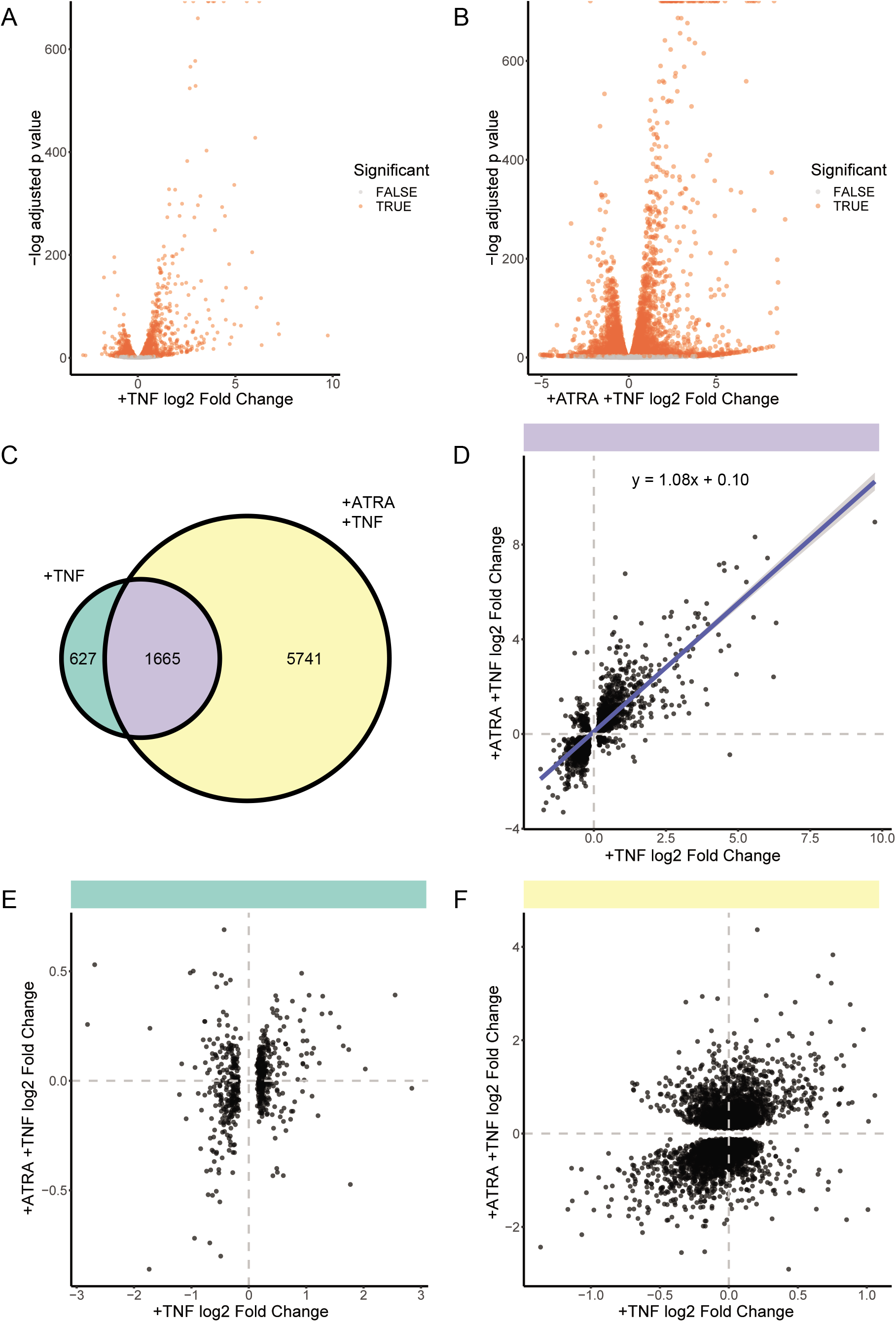
Transcriptional changes in promyelocytic and granulocytic forms of HL-60/S4 following two hours of TNF-α treatment. A) The Log2 fold change (log2FC) and adjusted p values of all analyzed genes in promyelocytes with and without TNF-α treatment. B) The log2FC and adjusted p values of all analyzed genes in granulocytes with and without TNF-α treatment. C) There is a shared and unique transcriptional response to TNF-α treatment of HL-60/S4 promyelocytes and granulocytes. D) The log2FC of genes that were significantly differentially expressed in both HL-60/S4 and HL-60/S4+ATRA cells were positively correlated (R2=0.65). However, there was a small subset of genes that were anticorrelated. E) The log2FC of genes that were only differentially expressed in HL-60/S4 cells did not correlate between HL-60/S4 and HL-60/S4+ATRA after TNF-α treatment (R2=0.04). E) The log2FC of genes that were only differentially expressed in HL-60/S4+ATRA cells did not correlate between HL-60/S4 and HL-60/S4+ATRA after TNF-α treatment (R2=0.15).

Genes that were significantly differentially expressed after TNF-α treatment in both the promyelocytic and granulocytic forms of HL-60/S4 cells exhibited strongly correlated log2 fold changes (p<2×10^−16^, R^2^=0.65). There was a small subset of differentially expressed genes that exhibited changes in opposite directions in promyelocytes and granulocytes (Fig 1D). Notably, the effect sizes of conserved gene expression changes were similar, with a slope of 1.08. If the observed differences in the numbers of differentially expressed (DE) genes were due to a different magnitude of effect, we would expect to see a correlation in fold change between genes that were differentially expressed in one cell type but not the other. However, the fold change of non-conserved genes did not strongly correlate between cell types (Fig 1E & F, R^2^=0.038, R^2^=0.15).

Collectively these results suggest that the difference in TNF-α responses is not simply due to granulocytic cells showing enhanced regulation of a conserved set of genes, resulting in a greater number that are significantly differentially expressed. Instead, additional sets of genes are changing transcriptional activity in TNF-α treated granulocytes.

### 2.2 Functional analysis of the TNF-α response in promyelocytes and granulocytes

Functional analysis of gene sets is affected by the available gene function information, and the method of enrichment analysis [40]. Therefore, we used multiple annotation sources (*i.e*. Molecular Signatures Databaste (MSigDB) hallmark gene sets [41], gene ontology (GO) [42], and Kyoto Encyclopedia of Genes and Genomes (KEGG) pathways [43]) to interrogate the TNF-α response in HL-60/S4 cells and identify robust enrichment results.

#### 2.2.1 Gene set enrichment analysis of promyelocytes and granulocytes treated with TNF

We first interrogated the global gene expression changes that occurred in the two different cell types after TNF-α treatment. Gene set enrichment analysis (GSEA) is a method that allows analysis of an entire differential expression dataset without using arbitrary significance thresholds [44]. Instead, the entire dataset is ranked by log2 fold change, and terms are considered enriched if they are overrepresented at the top or bottom of the ranked list. This approach allows for a differentiation between functional categories of genes that are upregulated or downregulated after treatment [45], and provides an overview of transcriptional changes.

We performed GSEA of hallmark gene sets (Supp table 3). We found a strong upregulation response to TNF-α, in both the promyelocytic and granulocytic forms of the HL-60/S4 cells, that was characterized by immune and cell death pathways (Fig 2). However, the granulocytic genes that were upregulated were also enriched for terms that included notch signaling and hypoxia. Transcripts associated with WNT and beta-catenin signaling were downregulated in promyelocytes (Fig 2).

**Figure 2.**
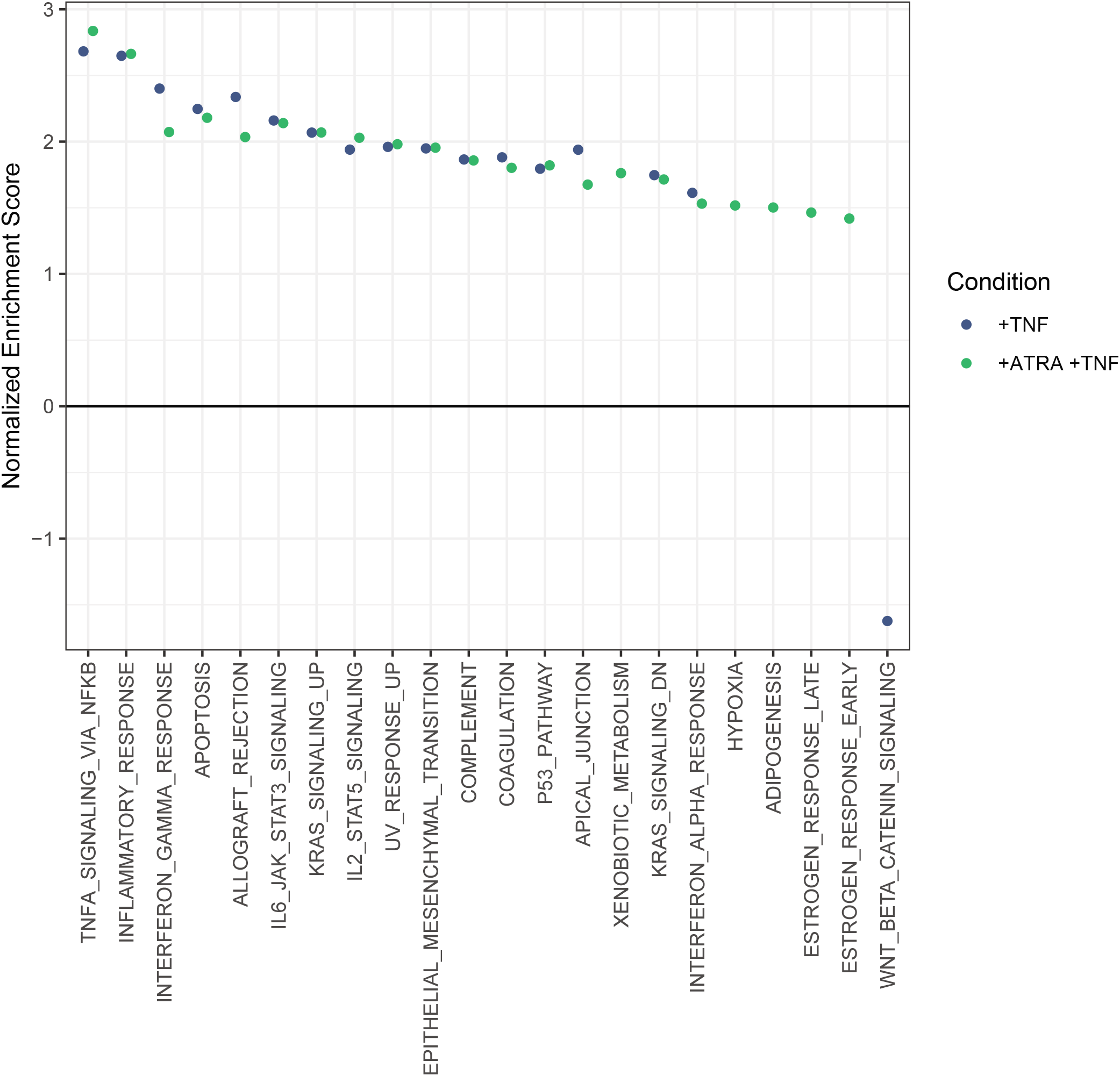
Transcriptional changes in promyelocytes and granulocytes treated with TNF. Gene set enrichment analysis (GSEA) of genes represented in the MSigDB Hallmark gene sets [1]. All represented genes were ranked by Log2FC, with no significance cutoff. The x axis shows significantly enriched gene sets (FDR<0.05). The normalized enrichment score (y axis) indicates whether a given gene set was overrepresented for transcripts that exhibited large fold changes. Predicted gene sets (e.g. TNFA signaling via NFKB, p53 pathway, and IFNγ response) were enriched in both conditions. No gene sets were enriched in HL-60/S4 but not HL-60/S4+ATRA. Six gene sets (adipogenesis, estrogen response early and late, hypoxia, IFNα response, and xenobiotic metabolism) were enriched in HL-60/S4+ATRA, but not HL-60/S4.

As observed in the differentially expressed genes, GSEA showed a strong conserved response with notable divergences between TNF-α stimulated promyelocytes and granulocytes.

#### 2.2.2 Functional analysis of conserved, cell type-specific, and opposite TNF-α responses

In order to further investigate the similarities and differences in the TNF-α response, we grouped the list of analyzed genes based on the direction and significance of the expression change in each cell type. Genes that were differentially expressed following TNF-α treatment were divided into four groups according to the nature of the correlation between the promyelocytic and granulocytic responses: 1) conserved; 2) promyelocyte-specific; 3) granulocyte-specific; and 3) opposite responses (Supp table 1). The first group represents the conserved response, *i.e*. the genes were significantly differentially expressed in the same direction in both promyelocytic and granulocytic forms of the HL-60/S4 cells. The next two groups represent cell type-specific responses – *i.e*. genes that were only differentially expressed in either promyelocytes or granulocytes in response to TNF-α treatment. The final group is comprised of the 157 genes that were significantly upregulated in the first cell type and significantly downregulated in the other cell type, or *vice versa*.

Having identified groups of genes that change in a conserved, cell-type specific, and opposite manner, we performed functional enrichment analyses to further investigate the responses to TNF-α in promyelocytes and granulocytes.

##### 2.2.2.1 A conserved TNF-α response

Genes that exhibited significant changes in the same direction in response to TNF-α treatment in both promyelocytic and granulocytic forms of the HL-60/S4 cells were more frequently upregulated than downregulated (~3:1). Notably, this trend was not maintained for cell type-specific gene expression changes (promyelocyte-specific ~1.2:1, granulocyte-specific ~0.9:1). There were nine enriched MSigDB hallmark gene sets [41] in the conserved response gene set (Fig 3A, Supp table 4), all of which are associated with well described effects of TNF-α stimulation; cytokine signaling, inflammation, and apoptosis. This was broadly consistent with the results of GSEA (Fig 2). The top three enriched pathways within KEGG were: NFKB Signaling; NOD-like receptor signaling; and TNF-α signaling pathways (Supp table 5). The top 10 enriched GO terms included interferon-gamma (IFNγ)-mediated signaling pathway, inflammatory response, and positive regulation of I-kappaB kinase/NFKB signaling (Supp table 6).

**Figure 3.**
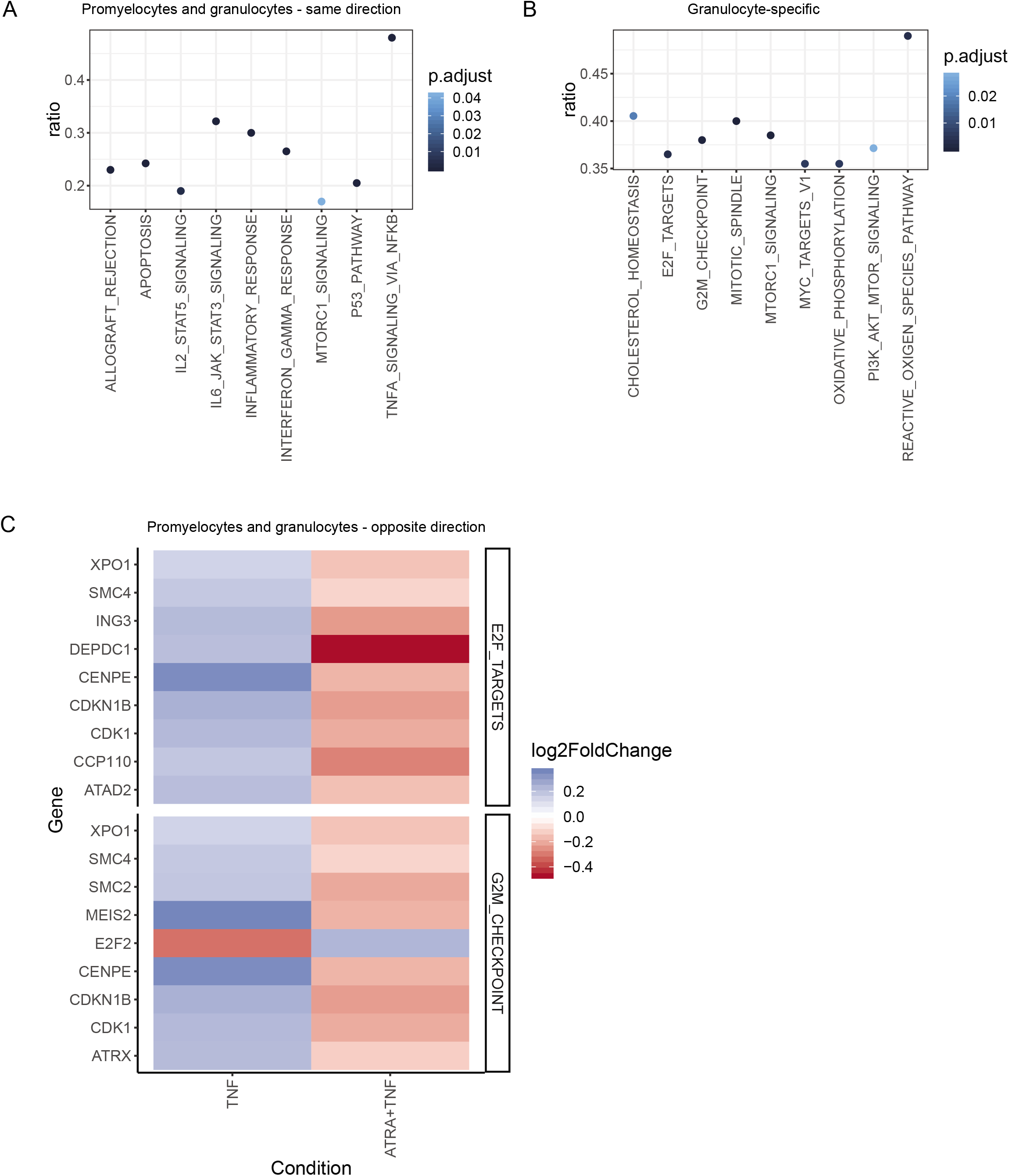
Gene set overrepresentation in differential expression subsets of TNF-α treated promyelocytes and granulocytes. A) Genes that were significantly differentially expressed in the same direction after TNF-α treatment were overrepresented in several gene sets canonically associated with TNF-α response. B) Genes that were only significantly differentially expressed in granulocytes were overrepresented in 9 gene sets, including 4 cell cycle associated gene sets. They were also overrepresented in the reactive oxygen species pathway, a neutrophilic response to TNF-α. C) Genes that were significantly differentially expressed in opposite directions were overrepresented in two cell-cycle associated gene sets. All differentially expressed hallmark G2M checkpoint genes were upregulated in promyelocytes, and downregulated in granulocytes cells, with the exception of E2F2, a transcription factor that promotes quiescence by binding to promoters and transcriptionally repressing cell cycle genes [2]. All differentially expressed E2F2 target genes were upregulated in promyelocytes, and downregulated in granulocytes.

In summary, using three different annotation databases and four different analysis approaches, we have shown that TNF-α treatment of both promyelocytic and granulocytic forms of HL-60/S4 cells induces a transcription profile associated with inflammatory responses and NFKB signaling.

##### 2.2.2.2 Granulocyte-specific TNF-α responses

The cell type-specific promyelocytic DE genes had no significant enrichments of MSigDB gene sets, KEGG pathways, or GO terms. Cell type-specific granulocytic DE genes were enriched for MSigDB sets related to cell cycle and energetics (Fig 3B, Supp table 7). The top five enriched KEGG pathways included Cell Cycle, Protein processing in endoplasmic reticulum, and Cellular senescence (Supp table 8). The six enriched GO terms included cell division, G2/M transition of mitotic cell cycle, protein poly- and de-ubiquitination, and neutrophil degranulation (Supp table 9). Thus, granulocytes, but not promyelocytes, exhibited transcriptional changes in genes involved in cell cycle and protein processing in response to TNF-α treatment.

##### 2.2.2.3 Opposite TNF-α responses in promyelocytes and granulocytes

Genes that were significantly differentially expressed in opposite directions in promyelocytes and granulocytes were enriched for G2M checkpoint and E2F targets within the MSigDB database (Fig 3C, Supp table 10). Differentially expressed genes in these sets were upregulated in promyelocytes and downregulated in granulocytes, with the exception of *E2F2*. *CDK1* is included in both of these cell-cycle associated gene sets, and is considered sufficient to drive the mammalian cell cycle [46]. As described above, the upregulation and downregulation of *CDK1* in promyelocytes and granulocytes respectively was confirmed by RT-qPCR (Supp table 2). The set of genes that was significantly expressed in opposite directions was also enriched for three terms, including mitotic chromosome condensation, within the GO database (Supp table 11). There was no enrichment for pathways within KEGG.

The inflammatory effects of TNF-α stimulation are shared by promyelocytes, but cell cycle repression and altered protein metabolism are TNF-α responses specific to granulocytes.

### 2.3 Granulocytic differentiation increases transcript levels of TNF and TNF receptors

We set out to investigate how differentiation into the granulocytic form could modulate the TNF-α response, and identify similarities in the transcriptional changes that occur during differentiation and acute TNF-α treatment. We were unable to analyze transcriptional changes that occurred during differentiation, as promyelocytic and granulocytic cells were treated with TNF-α in separate experiments. In order to assess gene expression changes associated with differentiation we analyzed publicly available RNA-seq data of HL-60/S4 cells differentiated into granulocytes (with ATRA) and macrophages (with TPA). Principal component analysis of variance stabilized transcripts confirmed clustering by differentiation status (Supp fig 1C).

Consistent with previous reports [30], this analysis confirmed that *TNF* expression was upregulated during differentiation into either granulocytic or macrophage forms of HL-60/S4 cells. The magnitude of *TNF* transcript upregulation was different in granulocytes (log_2_ fold change = 1.37) and macrophages (log_2_ fold change = 4.08) (Supp table 1). Differentiation also induced changes in the expression levels of the TNFRs. Consistent with previous observations [30], *TNFRSF1A* gene expression was increased (log_2_ fold change = 0.87) after ATRA treatment, but there was no significant change in expression following TPA treatment. By contrast, *TNFRSF1B* gene expression was increased in both conditions (log_2_ = 1.59 and 3.30 following ATRA and TPA treatment, respectively). This observed increase in mRNA expression of *TNF* and both TNF-α receptors may be one explanation for why granulocytes have an enhanced transcriptional response to TNF-α, compared to promyelocytes.

We investigated whether TNF-associated genes were enriched for transcriptional changes associated with differentiation into the granulocytic or macrophage form. We performed a GSEA of the transcriptional changes associated with differentiation into the granulocytic and macrophage form (Fig 4). As described previously, genes associated with cell-cycle terms (e.g. MYC targets and G2M checkpoint) were downregulated, while genes associated with inflammatory terms (e.g. IFNγ response, inflammatory response, and indeed TNF-α signaling via NFKB) were upregulated in both differentiation experiments (Fig 4).

**Figure 4.**
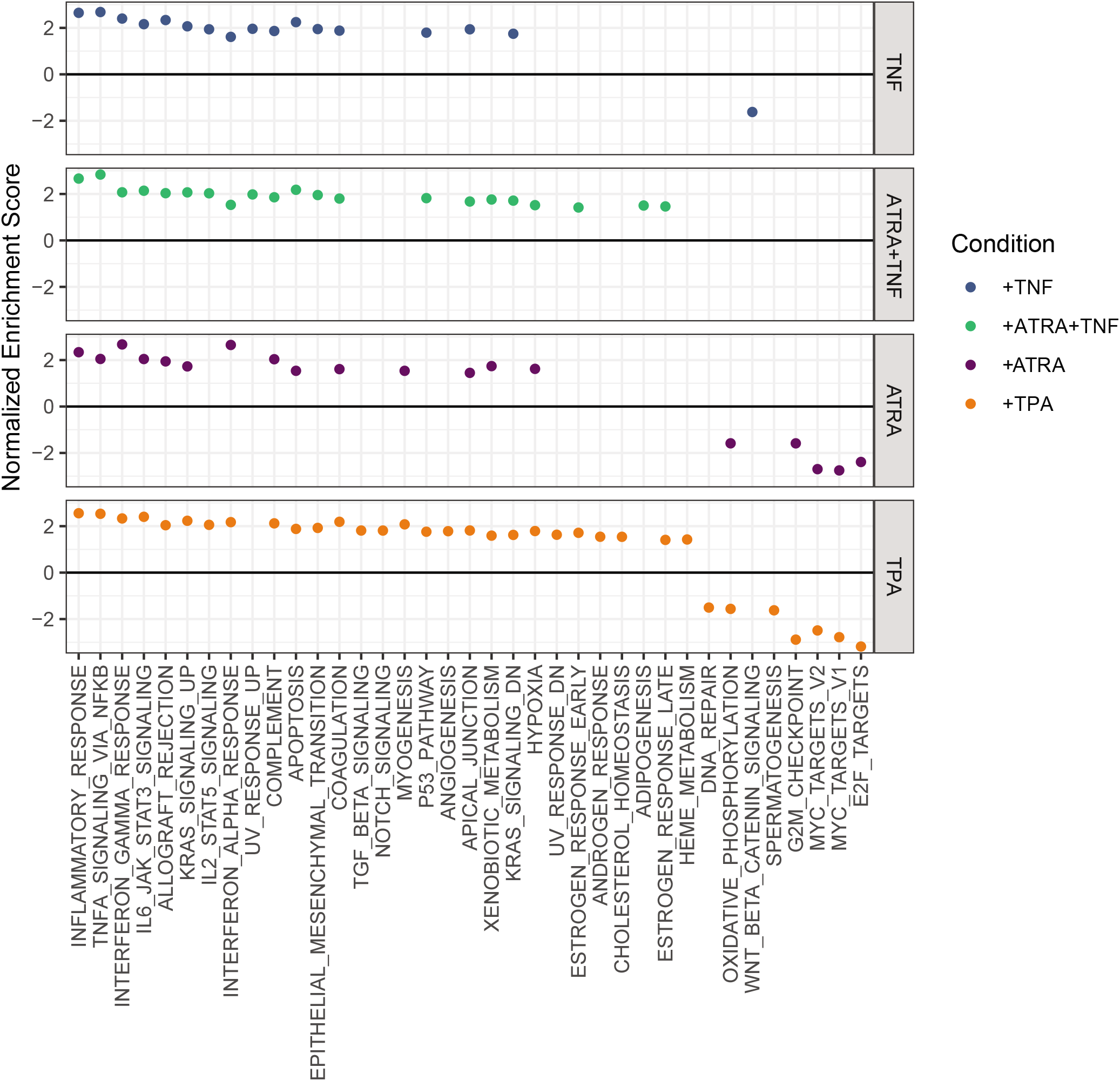
Transcriptional changes in promyelocytes differentiated into granulocytes with ATRA or macrophages with TPA, compared to TNF-α treatment of promyelocytes and granulocytes. Gene set enrichment analysis (GSEA) of genes represented in the MSigDB Hallmark gene sets [1]. All represented genes were ranked by Log2FC, with no significance cutoff. The x axis shows significantly enriched gene sets (FDR<0.05), and the y axis is calculated from the proportion of genes in the leading edge out of the number of ranked genes, and out of the gene set size. All conditions were associated with upregulated inflammatory gene sets, and both differentiation conditions were associated with downregulation of cell cycle gene sets.

As shown in Welch *et. al*. [30], differentiation into either the granulocytic or macrophage form resulted in higher inflammatory gene expression and lower cell cycle rate.

### 2.4 TNF-α treatment has more shared responses with TPA treatment than ATRA treatment

TNF-α treatment interacts with ATRA to augment differentiation of myeloid cells [33,34], but whether it enhances the differentiation into granulocytic cells or modifies the differentiation trajectory is unclear. We compared the transcriptional changes of TNF-α treated granulocytes with those associated with differentiation into granulocytes or macrophages.

Gene sets upregulated in TNF-α stimulated granulocytes and differentiated granulocytes or macrophages were enriched for ontological terms associated with hypoxia and xenobiotic metabolism. Notably, genes that were upregulated in promyelocytes treated with TNF-α did not show enrichment for these ontological terms. Notch signaling was upregulated only after TNF-treatment of promyelocytes or differentiation into macrophages. Intriguingly, both early and late estrogen response genes were upregulated in granulocytes treated with TNF-α, and macrophages, despite reports that TNF-α acts to oppose estrogen signaling in breast cancer [47]. Four terms were upregulated after TNF-α treatment of granulocytes and differentiation into macrophages: IL2 stat5 signaling, epithelial mesenchymal transition, apical junction, and KRAS signaling (genes downregulated by KRAS activation) (Fig 4).

To further compare the transcriptional changes that occur during differentiation into different cell types with the effects of TNF-α treatment, we assigned differentiation associated changes into four groups: 1) conserved; 2) granulocyte-specific; 3) macrophage-specific; and 4) opposite, *i.e*. significantly upregulated in granulocytes and significantly downregulated in macrophages, or vice versa (Supp table 1). The first group consists of genes that significantly change expression levels in the same direction after differentiation into either the granulocytic from with ATRA, or the macrophage form with TPA. The second and third groups contain genes that are differentially expressed only after differentiation into granulocytes or macrophages, but not both. The fourth group represents genes that are differentially expressed after differentiation into granulocytes and macrophages, but are upregulated in granulocytes and downregulated in macrophages, or vice versa. We analyzed these groups of genes for enrichment of MSigDB gene sets, KEGG pathways, and GO terms. GSEA identified the response that is conserved during differentiation into granulocytes or macrophages as being characterized by pro-inflammatory cytokine associated gene sets (*i.e*. IFNα, response, IFNγ response, TNF-α signaling via NFKB), and cell cycle associated gene sets (*i.e*. G2M checkpoint, MYC targets, and mitotic spindle) (Fig 5A). This finding was consistent with what was observed as being enriched within the KEGG pathways and GO terms (Supp table 13,14).

**Figure 5.**
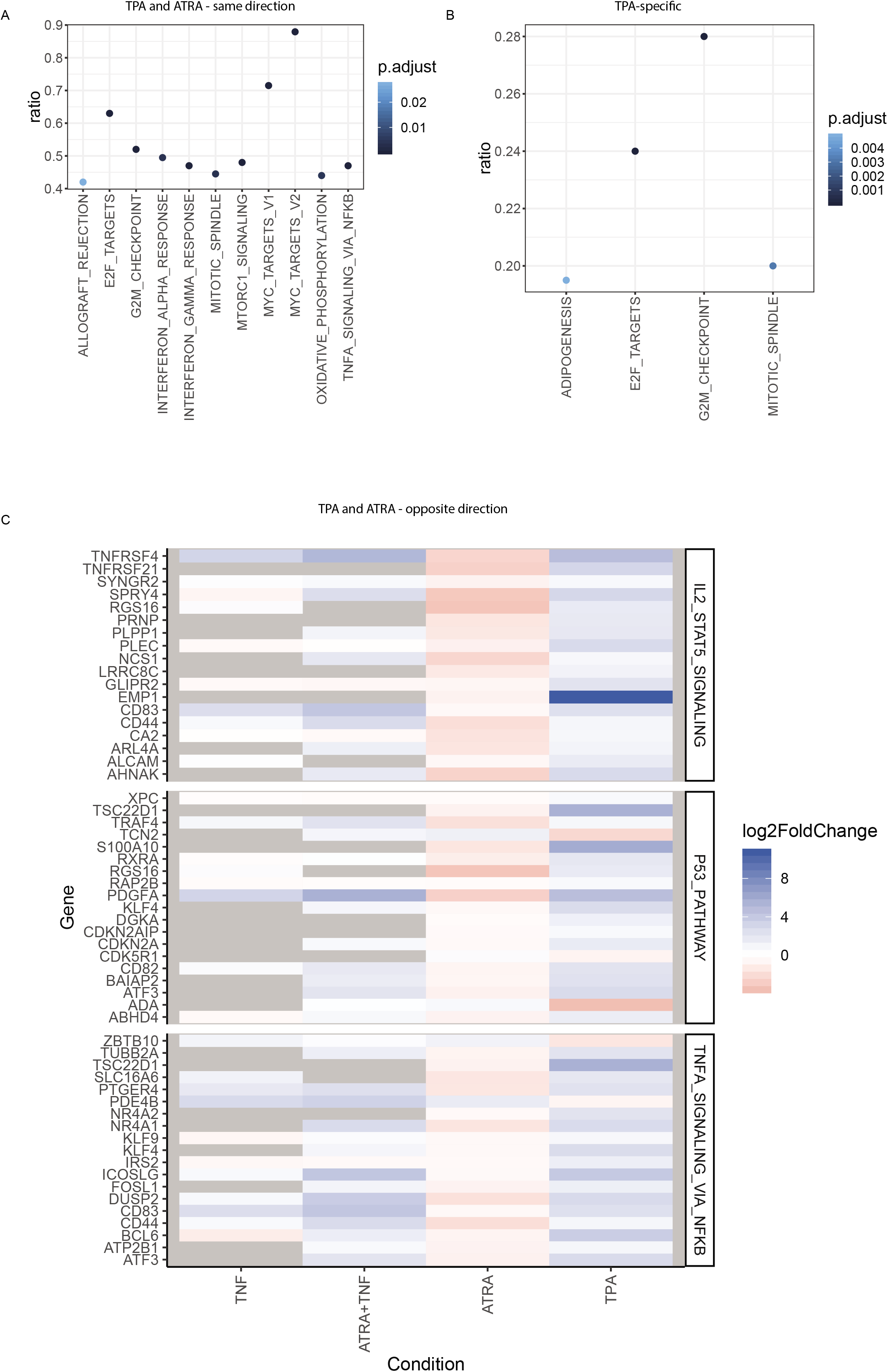
Gene set overrepresentation in differential expression subsets of promyelocytes differentiated into granulocytes or macrophages. A) Genes that were significantly differentially expressed in the same direction after differentiation were overrepresented in gene sets associated with inflammatory signaling and cell cycle. B) Genes that were only significantly differentially expressed in cells differentiated into macrophages with TPA were predominantly associated with cell cycle. C) Genes that were significantly differentially expressed in opposite directions were overrepresented in 3 gene sets: IL2 STAT5 signaling, p53 pathway, and TNF-α signaling via NFKB. A majority of genes in all categories increased expression after macrophage differentiation and in granulocytes treated with TNF-α, and decrease expression after granulocytic differentiation. Non-significant changes are indicated with grey.

Genes that changed expression after differentiation into granulocytes, but not macrophages, were not enriched for MSigDB gene sets or GO terms, but they were enriched for three KEGG pathways related to protein processing (Protein processing in endoplasmic reticulum, ubiquitin mediated proteolysis, and proteasome; Supp table 15). In contrast, genes that changed expression after differentiation into macrophages, but not granulocytes, were enriched for several MSigDB gene sets related to cell cycle (Fig 5B, Supp table 16). Enriched KEGG pathways included metabolism and protein processing (*i.e*. carbon metabolism and protein processing in endoplasmic reticulum), and the TNF-α signaling pathway (Supp table 17). Enriched GO terms included cell division, and terms related to RNA and protein regulation (*i.e*. regulation of transcription, DNA-templated, translational initiation).

#### 2.4.1 Convergence of transcriptional programs in macrophages and TNF-treated granulocytes

There are many similarities between the behaviors of HL-60/S4 cells differentiated into granulocytes and macrophages. However, there are also notable behavioral differences that include macrophages exhibiting a decreased cell cycle rate, increased survival, and adhesion to both surfaces and other cells [30].

We investigated the genes that were significantly differentially expressed with negatively correlated log2 fold changes after differentiation into granulocytes or macrophages –*i.e*. in opposite directions. This gene set was enriched for the IL2/STAT5 signaling, p53 pathway, and TNF-α signaling MSigDB hallmark terms (Fig 5C). 91% of these genes (102/112) increased expression after macrophage differentiation, and decreased expression after granulocytic differentiation. 65% of these genes (70/112) significantly increased expression in granulocytes treated with TNF-α, while only 10% (12/112) significantly decreased expression (Fig 5C). Six transcription factors (*ATF3, BCL6, FOSL1, KLF4, KLF9, NR4A1*) increased expression after macrophage differentiation and granulocytes treated with TNF-α, but decreased expression after granulocytic differentiation, which indicates a shared change in transcriptional programming. *CD83*, which is a marker of transdifferentiation of neutrophils into a dendritic-type cell, increased in both TNF-α treated cell types [48,49]. Given that macrophages and dendritic cells are phenotypically similar [50,51], *CD83* could be considered a marker of macrophage transdifferentiation.

Collectively our analyses suggest that TNF-α alters the transcriptional profile in HL-60/S4 cells consistent with a transition from a granulocytic to macrophage phenotype. Consistent with previous observations [33,34], TNF-α modifies the differentiation trajectory away from the ATRA induced granulocytic phenotype, towards a macrophage-like phenotype.

## Discussion

TNF-α treatment causes dramatic changes in the transcriptional programs of both promyelocytic and granulocytic HL-60/S4 cells. In this study we found that ATRA and TPA directed differentiation or TNF-α treatment of differentiated HL-60/S4 cells resulted in canonical TNF-α responses involving NKFB signaling, inflammatory signaling, p53 and apoptosis. Due to the reduced proliferation of granulocytic HL-60/S4 cells [30] we expected differential cell cycle effects of TNF-α treatment [24,28]. Previous work has shown increased proliferation in quiescent cells, while proliferating cells exhibited increased apoptosis after TNF-α treatment [24]. In contrast, we saw evidence of cell cycle repression in granulocytes, particularly at mitotic entry, while proliferating promyelocytes had an increase in cell cycle progression markers such as *CDK1* [46].

There are several factors that could explain the different responses of promyelocytes and granulocytes to TNF-α. Firstly, granulocytes have increased levels of endogenous TNF-α production. Not only does this increase the total TNF-α the cells are exposed to, but endogenously produced TNF-α is membrane-bound prior to processing [52]. The transmembrane and soluble forms of TNF-α have different effects [53], possibly due to the activation of different TNFRs. Membrane-bound TNF-α can activate both TNFR1 (*TNFRSF1A*) and TNFR2 (*TNFRSF1B*), but soluble TNF-α can only activate TNFR1 [54]. Granulocytic cells have higher levels of endogenous *TNF* expression (therefore, likely higher levels of transmembrane TNF-α) and concurrently higher levels of both *TNFRSF1A* and *TNFRSF1B* gene expression [55]. It has been previously proposed that increased levels of TNFR2 in granulocytic HL-60 cells explain their resistance to TNF-α induced apoptosis [35]. Thus, it may be not just the increased levels of receptors, but the ratio of TNFR1 to TNFR2 that determines the ultimate response to TNF-α.

The TNFR1 and TNFR2 receptors have uniquely stimulated pathways [56,57], however the two receptors also interact to produce a context specific TNF-α response [58,59]. Therefore, manipulating exogenous and endogenous levels of TNF-α or changing the ratio of the TNFR1 and TNFR2 receptors will provide greater insight into the phenotypic consequences of TNF-α treatment. Based on our data, we predict a dose dependent effect of total TNF-α on gene expression, whereby a low level of TNF-α is sufficient to induce regulation of the set of genes seen in the promyelocyte response, while higher levels of TNF-α are required to repress cell cycle. However, cell cycle repression likely also requires the presence of additional factors (*e.g*. TNFR-associated factors [60]) that are expressed during differentiation of the HL-60/S4 cells into the granulocytic form.

A reduction in the expression of cell cycle genes at the population level indicates a change in the proportion of cells at different stages of the cell cycle. Due to the massive transcriptional changes that occur during the progression through cell cycle [61–64], this may obscure other gene regulatory programs that are associated with alterations to cell function. Despite this limitation, we found evidence of a subset of TNF-α-regulated genes that were upregulated in TNF-treated granulocytes and following macrophage differentiation, but downregulated after granulocytic differentiation. These genes encoded a suite of transcription factors, and the dendritic cell surface marker CD83. Neutrophils that take on characteristics of antigen-presenting cells are often characterized by increased levels of CD83 [48,49], suggesting that TNF-α may be stimulating transdifferentiation. This is not unprecedented, as there is evidence that neutrophils have phenotypic plasticity [65], and that TNF-α can stimulate transdifferentiation [66–68]. However, further functional characterization of TNF-α-treated granulocytic HL-60/S4 cells is required to confirm this hypothesis.

A large-scale TNF-α response experiment treating many different cell types at various stages of differentiation would allow a network analysis of the transcriptional responses and annotation-free pathway discovery [69,70]. This would provide data to test our hypothesis that total TNF-α exposure correlates with repressive effects on the cell cycle. If this experiment were performed with single cell RNA-sequencing it would also enable the investigation of how individual TNF-α responses result in population-wide changes in cell cycle gene expression. For instance, do all cells respond to TNF-α stimulation with cytokine production, G2/M arrest, and apoptosis, or is there a heterogeneous response between different cells in a population? Moreover, single cell data would allow us to adjust for the cell cycle transcriptional effects, and identify differences that were previously masked by the population structure of unsynchronized cells.

Despite being one of the most highly studied genes in the human genome [71], the complex signaling [1] and context dependent effects [20,28,56] of TNF-α mean that much of its biology remains unknown. With the increasing accessibility of modern sequencing technologies and gene editing, high-throughput investigations of the TNF-α response and signaling will yield new results that enable the contextualization of TNF-α in cancer and immunology.

## Conclusion

HL-60/S4 cells show a conserved set of core responses to TNF-α treatment irrespective of the differentiation state of the cell. This is expected, since functional annotations represent canonical pathways and responses. However, granulocytes are more responsive to TNF-α, possibly due to priming from increased endogenous *TNF* expression, and increased levels of TNF-α receptors. Transcriptional changes indicate that TNF-α treatment represses cell cycle progression in granulocytes, but has the opposite effect on promyelocytes. This effect may be sensitive to the sum total of the exogenous and endogenous TNF-α levels. Finally, comparisons of transcriptional changes during differentiation and TNF-α treatment suggest that TNF-α treatment of granulocytes pushes them towards a macrophage transcriptional program. The context specific effects of TNF-α are likely mirrored in other models of innate immune signaling, and may contribute to the disparities seen between *in vitro* and *in vivo* studies of innate immune signaling.

## Methods

### Cell culture

HL-60/S4 cells (available from ATCC #CCL-3306) were cultured at 37°C, 5% CO_2_ in RPMI 1640 (ThermoFisher) supplemented with 10% fetal bovine serum (Moregate Biotech), 1% penicillin/streptomycin (ThermoFisher), 1% GlutaMAXTM (ThermoFisher) and 2mM CaCl_2_.

As required, cells were differentiated into a granulocytic phenotype with 1 μM all trans retinoic acid (ATRA) dissolved in ethanol (Sigma Aldrich) for four days as previously described ([30,36,72]). Cells were centrifuged (200xg, 5 min, room temp.) and suspended (undifferentiated = 1×10^6^ cells/mL; or differentiated = 5×10^5^ cells/mL) in fresh media (with 1μM ATRA for differentiated cells). Nuclear morphology and cell surface receptor changes during differentiation are available from Jacobson *et. al*. 2018 [36]. Hi-C and RNA-seq were used to identify translocations and other structural variants specific to HL-60/S4 [73]. Cells from the same seed-lot were used in all studies.

Cells (9 x 10^6^ per flask) were incubated at 37°C for 2 hours before addition of 16ng/mL TNF-α or vehicle in triplicate per condition. After 2 hours treatment, cells in each flask were lysed with TRIzol LS (Life Technologies), phase separated with chloroform, and RNA extracted with the Qiagen RNeasy mini kit (Qiagen, 74104). A summary of the conditions in this study, and the public data from Welch 2017 [30], are shown in table 1.

**Table 1.** Details of HL-60/S4 RNA-seq datasets. Four conditions in triplicate were generated in this study, while three conditions with four replicates were analysed from Welch et al. Cells were undifferentiated (promyelocyte), differentiated with ATRA (granulocyte) or TPA (macrophage). Promyelocytes and granulocytes in this study were treated with TNF or vehicle. The library preparation method in this study was ribosomal depletion to sequence all long RNAs, while Welch 2017 [30] used poly-A enrichment to sequence mRNA only. ‘Reads sequenced’ indicate the total number of paired-end reads generated per library. ‘Reads mapped’ indicate the number of read pairs aligned to the human genome with STAR [37]. ‘Reads assigned’ indicates the number of mapped reads assigned to genomic features (*e.g*. protein coding gene, non-coding RNA) with featureCounts [38].

| Data source | Library type | Differentiation agent | Cell type | Treatment | Replicate | Reads sequenced | Reads mapped | Reads assigned |
| --- | --- | --- | --- | --- | --- | --- | --- | --- |
| New data | Total | None | Promyelocyte | None | 1 | 51,259,672 | 48,256,979 | 31,775,702 |
|  |  |  |  |  | 2 | 45,409,486 | 43,088,708 | 26,931,181 |
|  |  |  |  |  | 3 | 47,722,497 | 45,321,199 | 28,412,920 |
| New data | Total | None | Promyelocyte | TNF | 1 | 45,401,741 | 42,548,543 | 26,121,834 |
|  |  |  |  |  | 2 | 46,622,711 | 44,003,082 | 28,053,460 |
|  |  |  |  |  | 3 | 52,842,736 | 50,124,478 | 31,895,061 |
| New data | Total | ATRA | Granulocyte | None | 1 | 46,965,601 | 44,427,313 | 24,592,785 |
|  |  |  |  |  | 2 | 53,329,425 | 50,564,062 | 28,655,822 |
|  |  |  |  |  | 3 | 52,544,954 | 49,873,305 | 28,687,969 |
| New data | Total | ATRA | Granulocyte | TNF | 1 | 44,130,932 | 41,753,860 | 23,800,117 |
|  |  |  |  |  | 2 | 55,591,032 | 52,524,970 | 29,318,607 |
|  |  |  |  |  | 3 | 44,662,624 | 42,149,334 | 22,829,189 |
| Welch 2017 | mRNA | None | Promyelocyte | None | 1 | 78,330,794 | 72,778,198 | 63,063,700 |
|  |  |  |  |  | 2 | 79,840,457 | 74,010,895 | 63,896,640 |
|  |  |  |  |  | 3 | 92,276,843 | 85,870,266 | 77,717,651 |
|  |  |  |  |  | 4 | 71,908,445 | 67,034,452 | 61,274,376 |
| Welch 2017 | mRNA | ATRA | Granulocyte | None | 1 | 78,215,359 | 73,037,649 | 63,738,330 |
|  |  |  |  |  | 2 | 79,385,831 | 73,007,606 | 65,788,452 |
|  |  |  |  |  | 3 | 67,293,855 | 61,810,327 | 55,209,128 |
|  |  |  |  |  | 4 | 74,792,543 | 68,660,444 | 61,293,657 |
| Welch 2017 | mRNA | TPA | Macrophage | None | 1 | 78,936,655 | 72,842,622 | 63,297,008 |
|  |  |  |  |  | 2 | 75,555,764 | 69,802,997 | 59,853,678 |
|  |  |  |  |  | 3 | 75,679,506 | 69,532,640 | 63,186,748 |
|  |  |  |  |  | 4 | 70,096,915 | 65,311,735 | 59,900,363 |

- 1 Mark Welch DB, Jauch A, Langowski J, et al. Transcriptomes reflect the phenotypes of undifferentiated, granulocyte and macrophage forms of HL-60/S4 cells. *Nucleus* 2017;8:222–237. - 2 Dobin A, Davis CA, Schlesinger F, et al. STAR: ultrafast universal RNA-seq aligner. *Bioinformatics* 2013;29:15–21. - 3 Liao Y, Smyth GK, Shi W. featureCounts: an efficient general purpose program for assigning sequence reads to genomic features. *Bioinformatics* 2014;30:923–930.

The full experiment was repeated and samples analyzed with RT-qPCR of *TNF, CDK1*, and *VCAM1* to validate the RNA-seq results (see below).

### RNA sequencing

Total RNA libraries were prepared by Annoroad Gene Technology Co., Ltd. (Beijing, China) using ribosomal RNA depletion with RiboZero Magnetic Gold Kit (Human/Mouse/Rat) and sequenced on an Illumina Hi-seq X 150PE (150 base pair paired end reads).

### Publicly available RNA-seq

mRNA-seq data of HL-60/S4 cells in three conditions in quadruplet [30] was downloaded from NCBI (http://www.ncbi.nlm.nih.gov/bioproject/303179). These conditions are: HL-60/S4 (promyelocytes), HL-60/S4 differentiated with ATRA (granulocytes), and HL-60/S4 differentiated with TPA (macrophage-like).

### RNA-seq analysis

Read quality was confirmed using FastQC v0.11.4. Paired end reads were aligned to hg38 and gencode annotations v27 using STAR [37] v2.5.3a with default settings. FeatureCounts [38] v1.5.2 was used to aggregate transcripts for gene-level analysis and quantify the reads with GENCODE [74] annotations v27. MultiQC [75] was used to summarize FastQC, STAR, and FeatureCounts outputs [75]. Mapping statistics are summarized in table 1.

Expressed genes were filtered (default settings) and differentially expressed genes (FDR<0.05) identified in DEseq2 [39] v1.16.1. Gene ontology enrichments were calculated with TopGo [42] v2.28.0 using the weight01 algorithm and the Fisher statistic. Categories containing <2 genes were removed, and p values adjusted for FDR. Kegg pathway analysis, GSEA, and enrichment analyses were performed using clusterProfiler [76]. Intersects were displayed with Vennerable [77], subsets extracted with dplyr [78], and plots were generated with ggplot2 [79] in R.

RNA-seq data was also used to check for mycoplasma contamination, as performed in [80]. All libraries were aligned to the genomes of four common mycoplasma species known to contaminate mammalian cell culture: *Mycoplasma hominis* ATCC 23114 (<u>NC_013511.1</u>), *M. hyorhinis* MCLD (<u>NC_017519.1</u>), *Mycoplasma fermentans* M64 (<u>NC_014921.1</u>) and *Acholeplasma laidlawii* PG-8A (<u>NC_010163.1</u>). The fasta genome sequences and genome annotation files were downloaded from NCBI Genome, and genome indices were created with STAR 2.6.0c. Due to their small genome size, the parameter genomeSAindexNbases was adjusted for each genome to the recommended log2(GenomeLength)/2 – 1. Zero reads from any of the 12 libraries aligned to any of the four mycoplasma genomes, confirming that our cultures were free from mycoplasma contamination.

### Quantitative real-time PCR

Total RNA was isolated using Trizol LS (Life Technologies). RNA was quantified using a NanoDrop spectrophotometer (NanoDrop Technologies). Isolated RNA was deoxyribonuclease I treated (Life Technologies). Single-stranded cDNA was synthesized from 1 μg of RNA using a High capacity cDNA RT kit (Thermofisher # 4368814), according to the manufacturer’s protocol. Real-time quantitative PCR analysis was carried out using predesigned PrimeTime^®^ Mini qPCR assays (Integrated DNA Technologies; Supp table 22) on a Lightcycler 480 (Roche). mRNA levels were normalized using reference genes (*GAPDH* and *COX4I1*). Log2 fold changes and errors was calculated using R using the delta-delta Ct method [81].

### Data availability

TNF-alpha treatment RNA-seq data is available on GEO, accession GSE120579. Differentiation RNA-seq data from [30] is publicly available on NCBI, http://www.ncbi.nlm.nih.gov/bioproject/303179. Analysis of processed RNA-seq data is available on github, https://github.com/jacel/TNF_HL60_S4.

## Additional files

**Additional file 1. Supplementary tables 1-22.** 1. Significantly differentially expressed genes in promyelocytes treated with TNF-a, differentiated into granulocytes and macrophages, and granulocytes treated with TNF-a. 2. RT-qPCR and RNA-seq of *CDK1, TNF*, and *VCAM1*. 3-21. Functional analysis summary tables of GSEA, GO, KEGG, and hallmark terms enriched in subsets of differentially expressed genes. 22. Primers and probes used for RT-qPCR validation.

**Additional file 2. Supplementary figure 1.** PCA of VST-normalized transcript counts show conditions cluster together in A) promyelocytes treated with TNF-α, B) granulocytes treated with TNF-α, and C) promyelocytes differentiated into granulocytes and macrophages.

## Authors’ Contributions

ECJ performed the experiments, analyzed and interpreted the data, and wrote the manuscript with support from JMO, JKP, and MHV. LJ performed and analyzed the RT-qPCR validation. All authors contributed to revisions and approved the final manuscript.

## Acknowledgements

The authors would like to thank Peter Shepherd and the members of the JMO lab group for comments and discussion.

