## Supplementary figure 1 for "TNF-α differentially regulates cell cycle genes in promyelocytic and granulocytic HL-60/S4 cells"

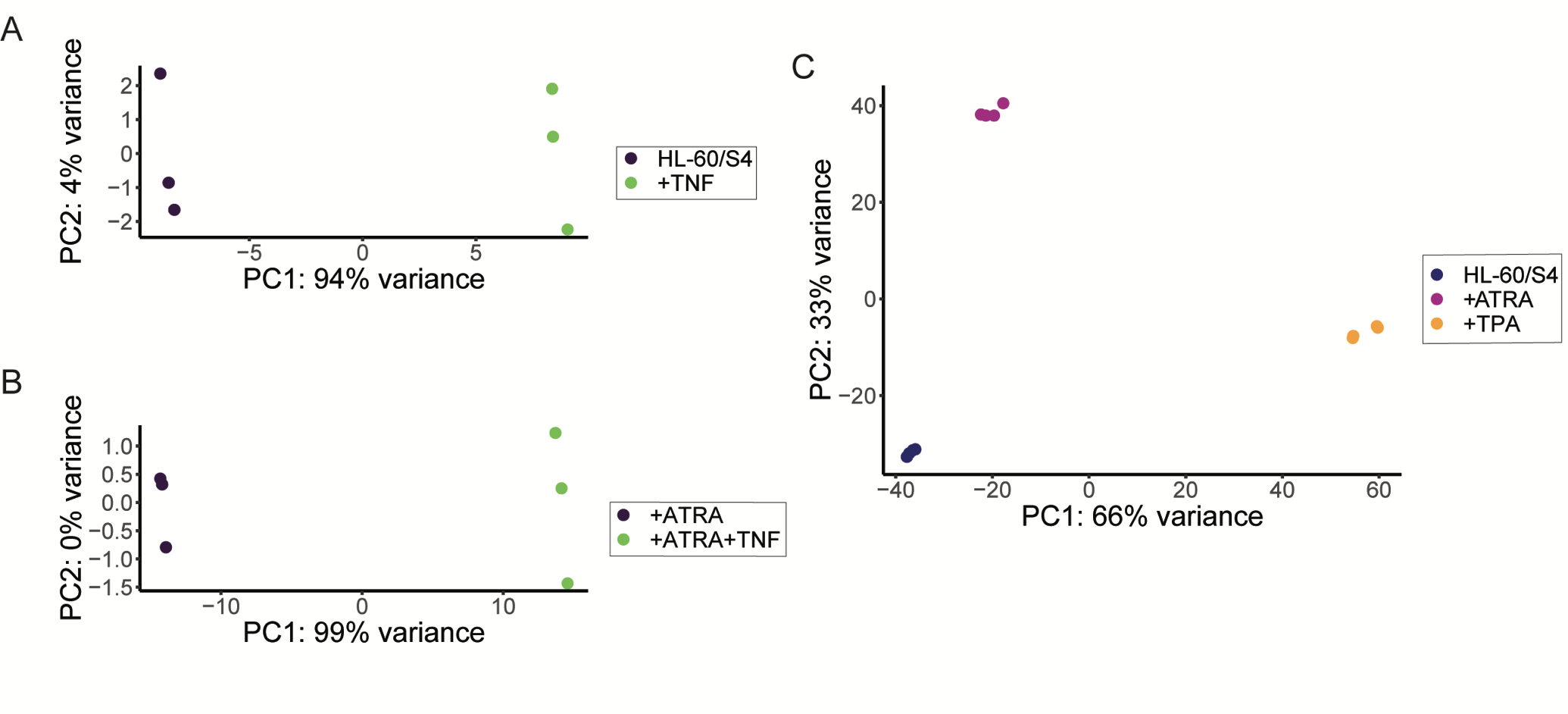

Supplementary Figure 1. Principle component analysis (PCA) of variance stabilised transcripts in HL60/S4 cells with A) TNF-α treatment for two hours; B) ATRA-differentiation for four days, then TNF-α treatment for two hours; C) Differentiation with ATRA, TPA, or vehicle for four days.
